# Tri-methylation of Histone H3 Lysine 4 Facilitates Gene Expression in Ageing Cells

**DOI:** 10.1101/238048

**Authors:** Cristina Cruz, Monica Della Rosa, Christel Krueger, Qian Gao, Lucy Field, Jonathan Houseley

**Affiliations:** Epigenetics Programme, The Babraham Institute, Babraham, Cambridge, UK; Present address: Adaptimmune Ltd, 60 Jubilee Avenue, Milton Park, Abingdon, Oxfordshire, UK; Present address: Molecular Haematology Unit, The MRC Weatherall Institute of Molecular Medicine, Oxford, UK

**Author notes:** These authors contributed equally to this research (Cristina Cruz, Monica Della Rosa).

## Abstract

Transcription of protein coding genes is accompanied by recruitment of COMPASS to promoter-proximal chromatin, which deposits di- and tri-methylation on histone H3 lysine 4 (H3K4) to form H3K4me2 and H3K4me3. Here we determine the importance of COMPASS in maintaining gene expression across lifespan in budding yeast. We find that COMPASS mutations dramatically reduce replicative lifespan and cause widespread gene expression defects. Known repressive functions of H3K4me2 are progressively lost with age, while hundreds of genes become dependent on H3K4me3 for full expression. Induction of these H3K4me3 dependent genes is also impacted in young cells lacking COMPASS components including the H3K4me3-specific factor Spp1. Remarkably, the genome-wide occurrence of H3K4me3 is progressively reduced with age despite widespread transcriptional induction, minimising the normal positive correlation between promoter H3K4me3 and gene expression. Our results provide clear evidence that H3K4me3 is required to attain normal expression levels of many genes across organismal lifespan.

## Introduction

H3K4me3 is ubiquitously observed on nucleosomes at the 5’ end of eukaryotic genes undergoing active transcription by RNA polymerase II (see for example (Barski et al., 2007; Bernstein et al., 2005; Liu et al., 2005; Zhang et al., 2009)). The tight correlation between promoter H3K4me3 and transcriptional activity has led to H3K4me3 being widely considered as an activating mark, however direct evidence for a role in steady-state transcription or gene induction is rare and controversial (reviewed in (Howe et al., 2017)).

H3K4me3 is primarily deposited by the highly conserved COMPASS complexes (Briggs et al., 2001; Krogan et al., 2002; Miller et al., 2001; Roguev et al., 2001; Santos-Rosa et al., 2002). Budding yeast has a single COMPASS complex containing the catalytic SET-domain protein Set1 and core structural proteins Swd1 and Swd3, along with Sdc1, Bre2, Swd2 and Spp1 (Figure 1A). Set1, Swd1 and Swd3 are required for all H3K4 methylation activity, while Sdc1 and Bre2 are necessary for di- and tri-methylation (Dehe et al., 2006; Schneider et al., 2005). Neither Swd2 nor Spp1 are required for H3K4 methylation *in vitro* (Takahashi et al., 2011), but mutation of Swd2 almost completely abrogates H3K4me3 and reduces H3K4me2 *in vivo* (Cheng et al., 2004; Lee et al., 2007), while loss of Spp1 dramatically reduces H3K4me3 *in vivo* without impacting H3K4me1/2 (Dehe et al., 2006; Schneider et al., 2005).

**Figure 1:**
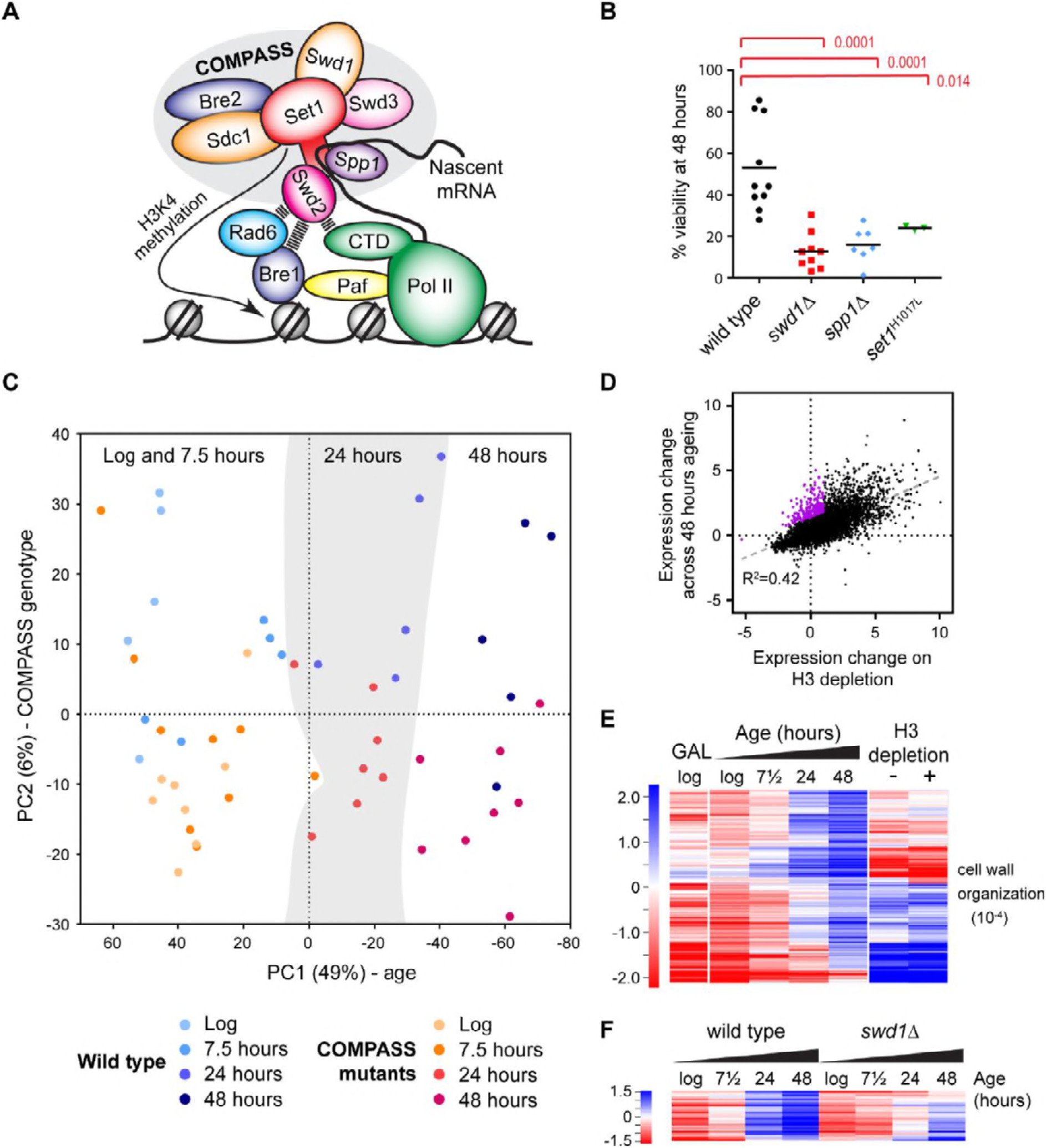
The ageing transcriptome in wild type and COMPASS mutants. **A**: Schematic of COMPASS recruitment for deposition of H3K4me3. **B**: Viability of mother cells after 48 hours in liquid YPD culture determined using the MEP system. p-values calculated by one-way ANOVA, n=10 (wild type), n=9 (swd1Δ), n=6 (spp1Δ), n=3 (set1^H1017L^). **C**: PCA plot of protein-coding gene expression for the 52 mRNAseq libraries generated from wild type and COMPASS mutants across 48 hours ageing. COMPASS mutants have been shown in the same colours for simplicity and to reflect their pooling in the initial DESeq2 analyses (Figure 2A,B). Vertical stripes show that wild type and COMPASS mutants are similarly distributed on PC1, which primarily reflects age. D: Plot of log2-transformed gene expression change from log phase to 48 hours ageing versus before to after H3 depletion. Genes that increase >2-fold more than average during ageing but increase <2-fold on H3 depletion are highlighted in purple. **E**: Hierarchical clustering analysis of log2-transformed protein-coding mRNA change for genes differentially expressed between log phase and 48 hours ageing in wild type but not on H3 depletion or between glucose and galactose media. GO enrichment assigned using GOrilla as in Figure 1 – figure supplement 1D. **F**: Hierarchical clustering analysis across ageing for wild type and *swd1*Δ of the subset of genes from E that are significantly differentially expressed between wild-type and *swd1Δ* cells at 48 hours (DESeq2 p<0.01 n=5 wild type, n=4 swd1Δ).

Promoter recruitment of COMPASS starts with the RNA pol II elongation factors Paf and FACT, which activate the H2B-ubiquitylation activity of Rad6/Bre1 (Pavri et al., 2006); this activity, and a non-mutated H2B^123K^ substrate, are vital for COMPASS recruitment and di-/tri-methylation (Dover et al., 2002; Krogan et al., 2003; Sun and Allis, 2002; Wood et al., 2003). Activation of Rad6/Bre1 and COMPASS recruitment also require interactions with the Ser-5 phosphorylated CTD of RNA polymerase II, providing a tight transcriptional connection (Krogan et al., 2003; Ng et al., 2003; Xiao et al., 2005). COMPASS recruitment occurs through interactions of Swd2 with active H2B-associated Rad6/Bre1, and of Set1/Spp1 with nascent mRNA (Battaglia et al., 2017; Luciano et al., 2017; Sayou et al., 2017; Thornton et al., 2014). In mammals, additional CxxC domains in Cfp1 (the orthologue of yeast Spp1) and some SET-domain proteins further specify and diversify COMPASS activity, allowing DNA methylation-dependent activity and selective recruitment of COMPASS to promoter and enhancer sequences (Brown et al., 2017; Clouaire et al., 2012; Hu et al., 2017; Thomson et al., 2010).

The importance of H3K4me3 in transcription has been difficult to study as loss of Set1 also abrogates H3K4me1 and H3K4me2. Even so, loss of Set1 does not drastically impair transcription in yeast, and the extent to which subsets of genes are mis-regulated in *set1Δ* cells has been controversial with later studies reporting mostly gene upregulation (Guillemette et al., 2011; Margaritis et al., 2012; Miller et al., 2001; Santos-Rosa et al., 2002; Venkatasubrahmanyam et al., 2007; Weiner et al., 2012). This suggests a primarily repressive role for H3K4 methylation, particularly evident for ribosome synthesis factors (Weiner et al., 2012), non-coding RNA-regulated genes (Berretta et al., 2008; Camblong et al., 2009; Pinskaya et al., 2009) and heterochromatic regions (Briggs et al., 2001; Li et al., 2006; Nislow et al., 1997; Venkatasubrahmanyam et al., 2007). To determine the importance of H3K4me3 specifically, studies in yeast have focused on cells lacking Spp1 which is required for most (but not all) H3K4me3, and have observed virtually no impact on steady state gene expression or on acute gene induction during media shifts (Howe et al., 2017; Margaritis et al., 2012). In contrast, loss of CFP1 in mammalian cells has concrete effects on gene expression, causing both up- and down-regulation of many genes in mouse ES cells and reduced transcription in oocytes (Brown et al., 2017; Yu et al., 2017). Interpretation of these observations is complicated by the additional binding modes of CFP1 that target COMPASS to promoters and enhancers, so loss of H3K4me3 may not be the only effect of CFP1 depletion (Brown et al., 2017; Hu et al., 2017). One specific report ties H3K4me3 mechanistically to activation of p53-regulated genes in response to DNA damage through recruitment of TAF3 in HCT116 cells, however this may be cell-type specific as the same response was not impaired in mouse ES cells lacking CFP1 (Clouaire et al., 2014; Lauberth et al., 2013). Overall, H3K4 methylation has the potential to both repress and activate transcription, but a general role for H3K4me3 in transcriptional activation remains unproven.

An association between histone modifications and ageing is well recognised (reviewed in (Maleszewska et al., 2016)). For example, defects in H3K4, H3K36 and H3K27 methylation all affect lifespan presumably through impacts on gene expression, though in most cases the downstream expression differences remain obscure (Alvares et al., 2014; Greer et al., 2010; Li et al., 2010; Maures et al., 2011; McColl et al., 2008; Sen et al., 2015; Siebold et al., 2010). Remarkably, in yeast and cultured human cells ageing is accompanied by a gross genome-wide loss of histones (Feser et al., 2010; Ivanov et al., 2013; O’Sullivan et al., 2010), leading to an opening of chromatin structure particularly at repressed promoters and causing widespread induction of normally repressed genes (Hu et al., 2014). General opening of chromatin structure has therefore been proposed to underlie many age-linked gene expression differences (Hu et al., 2014), and it is easy to imagine how alterations in canonical transcription-associated histone modifications could exacerbate these phenotypes. However, the impacts of histone modification defects on particular genes as cells age remain largely unknown due to the difficulty of de-convolving these from the massive impact of the ageing process itself.

Here we analyse the importance of H3K4me3 in facilitating gene expression across the lifetime of budding yeast. We find that the tight correlation between H3K4me3 and gene expression is progressively lost during ageing, but surprisingly that H3K4me3 is critical for the full expression of many genes induced with age. We directly validate the importance of H3K4 trimethylation in the expression of a subset of genes, demonstrating a direct role for H3K4me3 in maintaining normal expression of many genes across organismal lifespan.

## Results

### A transcriptomic dataset for ageing wild type and COMPASS mutants

A published micromanipulation screen of 264 yeast mutants found that cells lacking COMPASS components Swd1 or Swd3 have lifespans ~20% shorter than wild type (Smith et al., 2008). To determine the importance of COMPASS in controlling gene expression and chromatin structure across replicative lifespan we introduced selected COMPASS mutants into the Mother Enrichment Program (MEP) background; the MEP facilitates isolation of highly aged yeast in sufficient quantities for standard molecular techniques, and provides simple comparative methods to assess lifespan (Lindstrom and Gottschling, 2009). The lifespan methods, although less precise than microdissection or microfluidics, are performed in the same growth conditions as for other molecular techniques and therefore allow integration of lifespan and molecular data. As expected, MEP *swd1Δ* cells show a substantial and significant defect in replicative lifespan, being only ~50% viable at 24 hr and almost completely inviable after 48 hr in YPD (Figure 1 – figure supplement 1A). Importantly, this lifespan defect was recapitulated in catalytic dead *set1^HW17L^* and H3K4me3-defective spp1Δ mutants, showing that the lifespan defect results specifically from loss of H3K4me3 (Figure 1B and Figure 1 – figure supplement 1B). COMPASS mutants also have a reduced chronological lifespan due to stimulation of apoptosis by the H3K79 methyltransferase Dot1 (Walter et al., 2014); however no suppression of the replicative lifespan defect was observed in *swd1Δ dot1Δ* double mutants (Figure 1 – figure supplement 1C) so the replicative and chronological lifespan defects cannot be attributed to the same mechanism. Therefore, although *spp1Δ* mutants have a very weak phenotype during normal growth, this mutation dramatically impairs lifespan indicating that H3K4me3 has a prominent function in ageing cells.

To understand the gene expression differences between wild-type and COMPASS mutants, we obtained mRNAseq profiles from multiple replicates of wild-type, *swd1Δ*, *set1^H1017L^* and *spp1Δ* cells in log phase and aged for 7.5, 24 or 48 hours in YPD, forming a dataset of 52 libraries (Table 1). Principle component (PC) analysis of this dataset shows that PC1, 49% of variance, corresponds to age with both wild type and COMPASS mutant samples of equivalent ages having similar PC1 values (Figure 1C vertical stripes), although the log and 7.5 hour samples are similar and do not segregate well. PC2, 6% of variance, broadly segregates wild type from COMPASS mutant libraries (Figure 1C, note that wild type samples in blue lie towards the top whilst COMPASS mutants in yellow/red are largely in the lower half). The small effect of COMPASS genotype is not unexpected given previous reports, but shows that the gene expression phenotype caused by COMPASS mutation is separable from the gene expression phenotype caused by ageing. This gives us confidence that gene expression differences at equivalent times do not simply represent faster or slower ageing, which may otherwise be suggested based on the truncated lifespan of the COMPASS mutants.

**Table 1:**
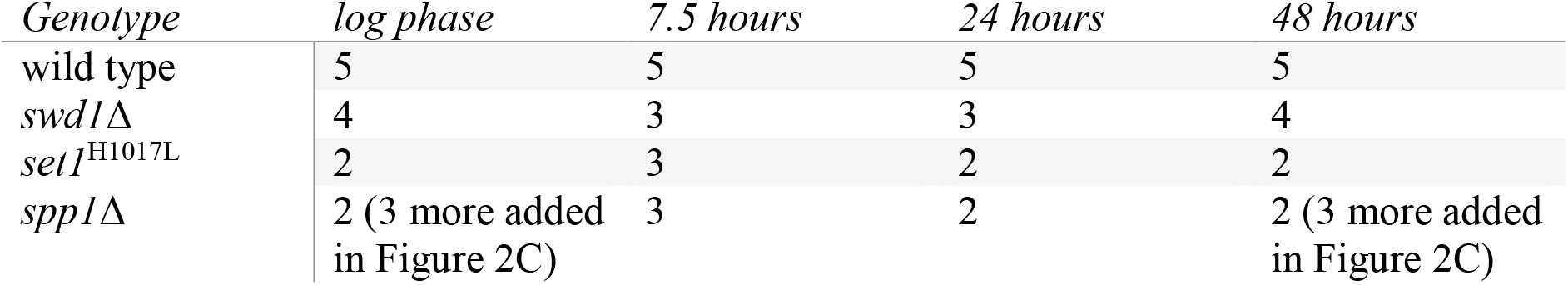
mRNAseq libraries analysed in this work

Differential expression analysis of the wild-type dataset identified 2842 out of 6662 annotated open reading frames (ORFs) that differ significantly between log and 48 hrs, with very similar results in the *swd1Δ* mutant (Figure 1 – figure supplement 1D). GO analysis of these ORFs confirmed many previous findings: translation-associated terms are down-regulated relative to average with age (Hu et al., 2014; Janssens et al., 2015; Kamei et al., 2014; Yiu et al., 2008), while *cell wall organisation, hexose transport, sporulation, tricarboxylic acid cycle* and *DNA integration* (i.e.: transposon activity) are upregulated as variously reported (Hu et al., 2014; Kamei et al., 2014; Koc et al., 2004; Lesur and Campbell, 2004). Genes upregulated with age are generally expressed at low levels in young cells, while genes that are highly expressed in young cells tend to be down-regulated with age relative to average as previously observed (Figure 1 – figure supplement 2)(Hu et al., 2014); in absolute terms, it has been shown that all yeast genes are actually induced to a greater or lesser extent during ageing, and we therefore refer to all gene expression changes as relative to average (Hu et al., 2014).

Age-related gene induction has been directly attributed to loss of histones, and we observe a strong correlation between age-linked gene expression and previously described changes following histone H3 depletion (Figure 1D)(Gossett and Lieb, 2012; Hu et al., 2014). We were interested to know if any particular category of genes is upregulated with age but not histone depletion, and so filtered for genes that are upregulated 2-fold more than average with age but increase less than 2-fold on H3 depletion (Figure 1D purple). We also filtered out genes repressed by the galactose to glucose shift used for H3 depletion in the Gossett and Lieb dataset, as the effect of H3 depletion for these genes is not determined. This left a core set of 204 genes, enriched for *cell wall organization* functions, that are robustly upregulated during ageing but not on H3 depletion (Figure 1E). This demonstrates that candidate age-linked gene expression programmes can be identified in yeast. Importantly 13% of these genes are significantly underexpressed in the *swd1Δ* mutant at 48 hours (Figure 1F), showing that loss of H3K4 methylation has a measureable impact on gene expression in aged cells.

### Requirement for H3K4me3 to facilitate age-linked gene induction

As a subset of ORFs are not properly induced in aged *swd1Δ* cells, we performed a more detailed differential expression analysis to identify H3K4 methylation-dependent genes. Compared to gene expression changes that occur during ageing the differences between wild type and COMPASS mutants are not large (Figure 1C), so we pooled samples in the initial analysis to increase statistical power. Given that all the tested COMPASS mutants reduced lifespan and may therefore have similar effects, we pooled together the replicate datasets for *swd1Δ*, set1^H1017L^ and *spp1Δ* for comparison to wild type (giving 7-9 COMPASS mutant replicates per time point, compared to 5 wild-type replicates). 107 genes were significantly different between wild type and COMPASS mutants at log phase, with more being differentially expressed at 24 and 48 hours (389 and 488), but few at 7.5 hours (39). Interestingly, log phase cells contain a subset of genes that are substantially over-expressed in COMPASS mutants (on average 3-fold), but this difference disappears with age (Figure 2A, top). In contrast, significantly under-expressed genes at log phase are on average only <1.5-fold lower than in wild type, but with age many more genes become under-expressed in COMPASS mutants and to a greater degree (2-fold on average, but many are 3-4-fold reduced) (Figure 2A, bottom). Genes that are over-expressed at log phase in COMPASS mutants (specifically *swd1Δ* and set1^H1017L^) remain over-expressed as ageing progresses but to a decreasing extent, whereas genes that are under-expressed in log phase remain under-expressed throughout ageing (Figure 2 – figure supplement 1A). These log phase data are in accord with previous results (Margaritis et al., 2012), but it is clear that the importance of COMPASS for gene expression changes with increasing age; in general, the repressive function of COMPASS has less impact and a positive function in gene expression is unmasked.

**Figure 2:**
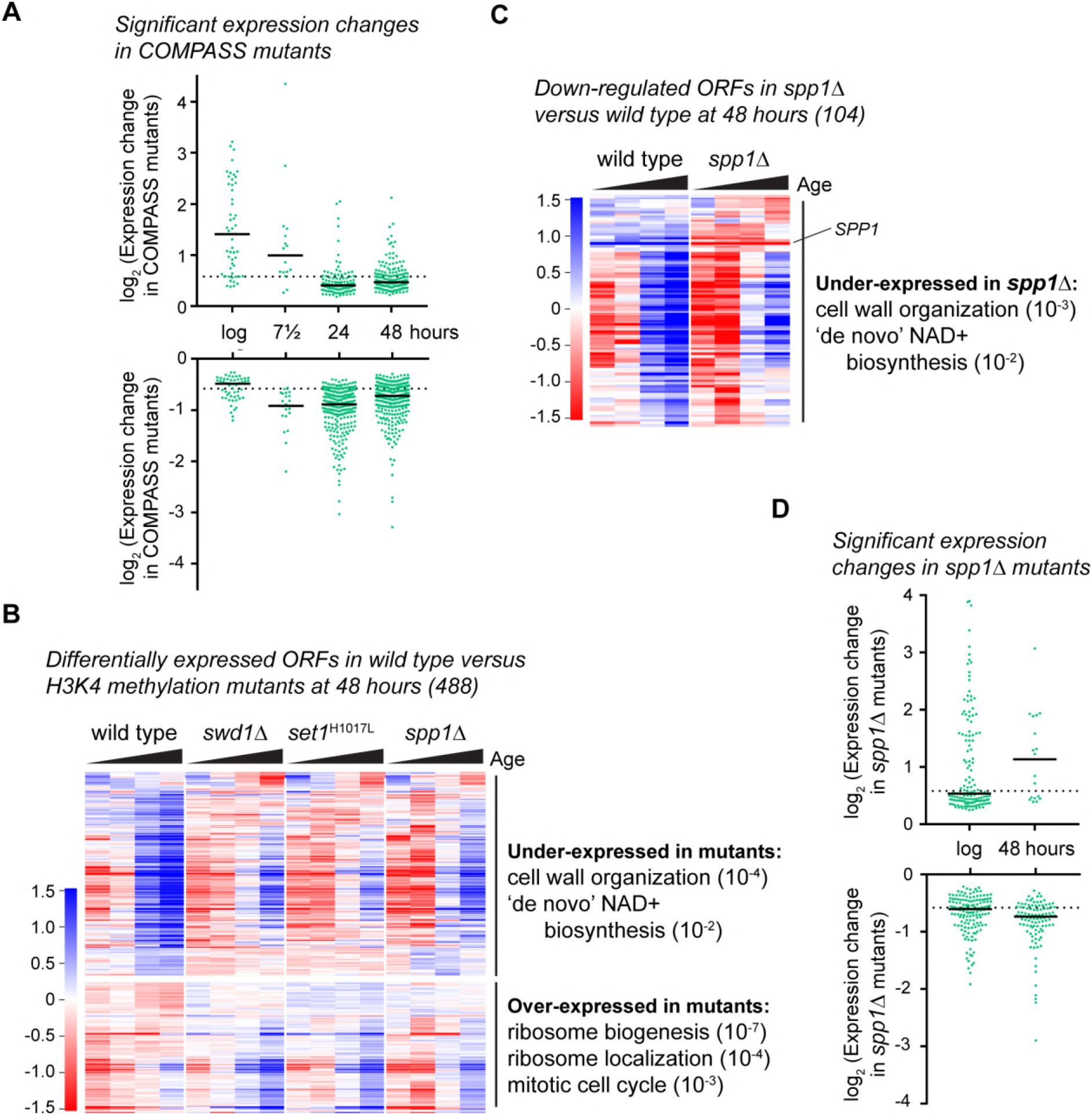
COMPASS is required for gene expression in ageing cells. **A**: Plot of log2-transformed read-count differences for significantly differentially expressed genes between wild type and COMPASS mutants at each ageing time point. Assessed using DEseq2 p<0.01, wild-type n=5 at each time point, COMPASS mutants n=8 at log, n=9 at 7.5 hr, n=7 at 24 hr and n=8 at 48 hr. Horizontal bars show median, dotted line indicates 1.5-fold change. **B**: Hierarchical clustering analysis of log2-transformed protein-coding mRNA change for genes significantly differentially expressed between wild-type and COMPASS mutants at 48 hours ageing, based on analysis in A. Separate time-courses are given for wild-type and each COMPASS mutant to show similar behaviour, GO analysis as for Figure 1E. **C**: Hierarchical clustering analysis of log2-transformed protein-coding mRNA change with age for genes significantly differentially expressed between wild-type and *spp1Δ* mutants at 48 hours ageing, assessed using DESeq2 p<0.05 n=5 per strain. GO analysis as Figure 1E. **D**: Distribution of significantly differentially expressed genes between wild type and *spp1Δ* at log and 48 hr, analysis as in A, n=5 for each set.

Starting with the 48 hour time point, we first confirmed that gene expression differences were consistent across replicates (Figure 2 – figure supplement 1B), then segregated under- and over-expressed genes and performed GO analyses (Figure 2B). Over-expressed genes (191) were primarily enriched for ribosome-related terms as expected (Weiner et al., 2012), but under-expressed genes (297) were enriched for *cell wall organisation* and *de novo NAD+ biosynthesis*, both of which are known to be important for maintenance of replicative lifespan (Bonkowski and Sinclair, 2016; Molon et al., 2017). Notably, the majority of genes affected by loss of COMPASS failed to be properly upregulated with age; this behaviour has not been observed in other transitions - Margaritis *et al.* examined the massive transcriptional reprogramming that accompanies the transition from stationary phase to log phase growth and found 220 genes mis-regulated of which only 24 (10%) were under-expressed in COMPASS mutants (Margaritis et al., 2012), compared to 297 (61%) of significantly altered genes that we observe in aged cells.

Differential expression in the pooled dataset could be attributed to mono-, di- or trimethylation of H3K4. To discover effects stemming purely from trimethylation, we sequenced three additional *spp1Δ* samples at log phase and after 48 hours ageing, giving five replicates in total. We employed a more relaxed significance threshold (p<0.05) for differential expression analysis to reflect the fact that *spp1Δ* is not a complete loss-of-function mutant and is therefore expected to cause a weaker phenotype (figure 1 – figure supplement 1B); this yielded 104 significantly under-expressed genes in aged *spp1Δ* cells compared to wild type, along with 19 over-expressed genes (Figure 2C). Despite the relaxed significance threshold, these differences remained highly consistent across replicates (Figure 2 – figure supplement 1C). GO analysis of the under-expressed genes again revealed significant enrichments for *cell wall biogenesis* and *de novo NAD+ biosynthesis* just as in the combined COMPASS mutant set, showing that proper expression of these genes is dependent on H3K4me3 (Figure 2C). Surprisingly, we also detected almost 300 genes differentially expressed between wild-type and *spp1Δ* log-phase cells, although the bulk of these showed very small differences (<1.5-fold) in contrast to the 48 hour samples, in which most differentially expressed genes were under-expressed and to a larger extent (>1.5 fold) (Figure 2D).

Overall, although COMPASS mutants are considered to have little impact on gene expression in log phase, we detect many genes (~4.5%) that require COMPASS to attain normal expression over the period of the yeast replicative lifespan.

### Importance of H3K4me3 in NAD+ biosynthesis gene induction

The failure of many genes to attain full expression during ageing in H3K4 methylation mutants indicates a role for this mark in facilitating gene expression. However, the complexity of the ageing process and the short lifespan of COMPASS mutants may render such effects indirect. To validate this association we focused on the *BNA* genes, which are regulated in a fairly simple manner by promoter binding of the repressive NAD+-dependent histone deacetylase Hst1 (Figure 3A)(Bedalov et al., 2003). *BNA1, BNA2* and *BNA4-7* encode the enzymes required for biosynthesis of the NAD+-precursor NaMN from tryptophan (Figure 3A), and differential expression of these genes underlies the GO enrichment for *de novo biosynthesis of NAD+* in ageing COMPASS mutants described in the previous section.

**Figure 3:**
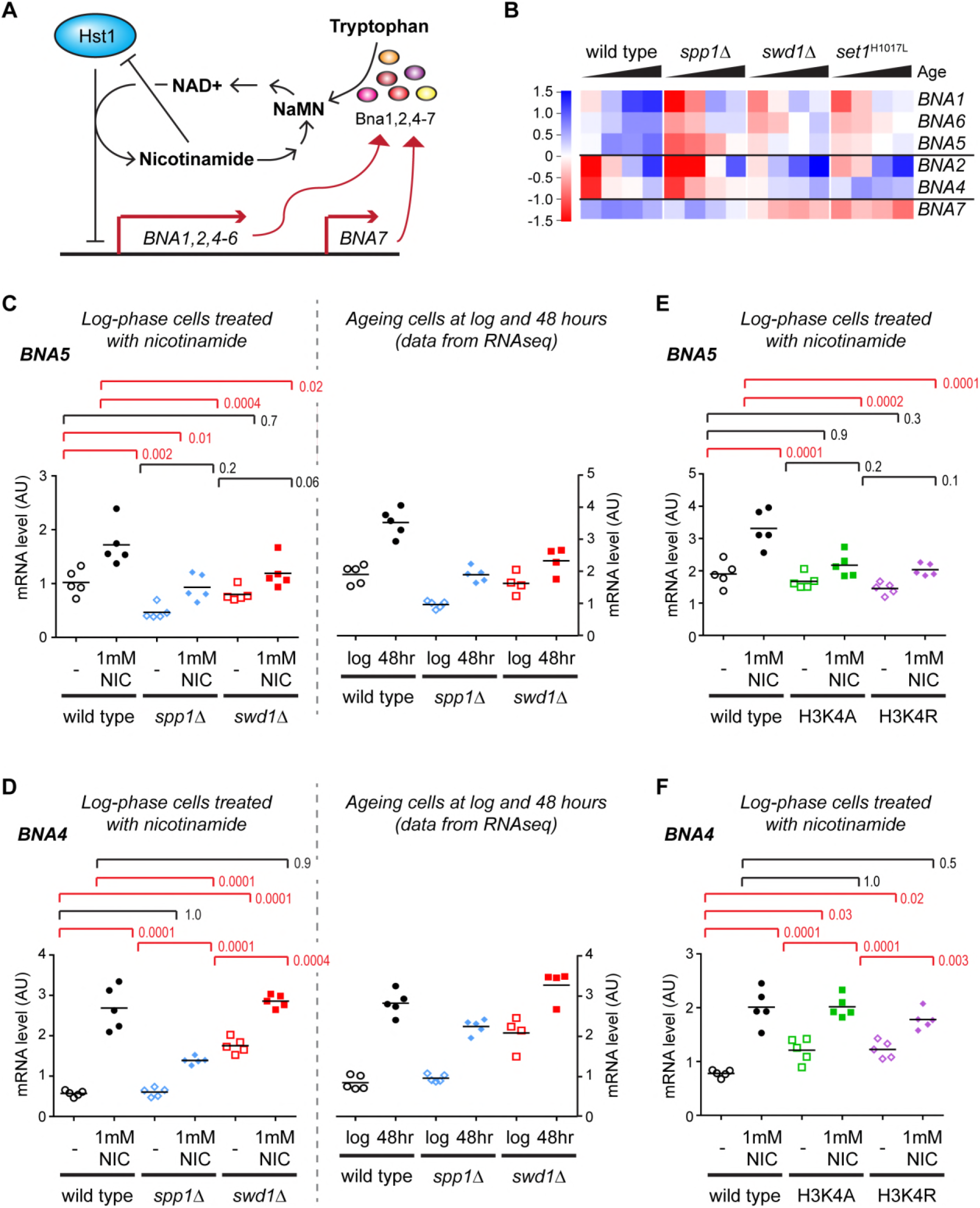
Validation of gene expression differences in COMPASS mutants. **A**: Schematic representation of the regulatory feedback system controlling expression of the *BNA* genes which encode enzymes in the pathway for NAD+ biosynthesis from tryptophan via nicotinic acid mononucleotide (NaMN). Essentially, Hst1 uses NAD+ as a cofactor to repress the *BNA* genes, so that when NAD+ is low the expression of the *BNA* genes rises to increase NAD+ biosynthesis. NAD+ is converted to nicotinamide by Hst1, which is a product inhibitor and limits the repressive activity. Exogenous nicotinamide is efficiently taken up by cells, allowing Hst1 inhibition in culture. NAD+ can also be synthesised from nicotinamide riboside, but this does not involve the BNA genes. **B**: Hierarchical clustering analysis of log2-transformed protein-coding mRNA change for genes in the tryptophan to NaMN biosynthesis pathway in wild type and COMPASS mutants across ageing. **C**: Northern blot quantification of *BNA5* mRNA level relative to ribosomal RNA in log phase BY4741 haploid cells grown in YPD and treated for 6 hours ±1 mM nicotinamide (left panel) compared to mRNA abundance measurements for *BNA5* derived from sequencing data for log and 48 hour aged cells (right panel). p-values were calculated by one-way ANOVA, significant differences are highlighted in red, n=5 for each category. **D**: Analysis of *BNA4* expression as in C. **E**: Analysis of *BNA5* expression in histone point mutants ± nicotinamide, performed as in C. **F**: Analysis of *BNA4* expression in histone point mutants ± nicotinamide, performed as in C.

We first assessed the expression of these genes individually across age in the RNAseq dataset. *BNA1, BNA2, BNA4, BNA5* and *BNA6* are all induced with age and this induction is impaired in *spp1* Δ, with *BNA7* forming an outlier that is unchanged by age or *SPP1* deletion (Figure 3B compare wild type to spp1Δ). The effect of *swd1Δ* and set1^H1017L^ is more complex: *BNA1, BNA5* and *BNA6* also show a defect in aged-linked induction in these mutants; *BNA2* and *BNA4* induce less but from a higher basal level; *BNA7* shows reduced basal expression. These data reveal that loss of H3K4me3 has a different effect on the *BNA* genes to loss of all H3K4 methylation.

Usefully, the direct negative regulation of the *BNA* genes by Hst1 allowed us to quantify induction in non-ageing log-phase cells through treatment with the Hst1 inhibitor nicotinamide (Figure 3A, Hst1 is subject to product inhibition by nicotinamide). As ageing was not required, we performed these experiments in standard BY4741 haploid cells without the substantial genetic modifications required for the MEP system. Five replicate sets of wild-type, *spp1Δ* and *swd1Δ* cells were grown in YPD, treated for 6 hours ±1 mM nicotinamide, RNA was extracted and expression of *BNA1* and *BNA4-7* determined in treated and untreated cells relative to ribosomal RNA by northern blot.

The induced gene expression level of *BNA1, BNA4* and *BNA5* was significantly reduced in *spp1Δ* cells, with *BNA6* showing a similar trend (Figure 3C,D and Figure 3 – figure supplement 1A,B). Furthermore *BNA1, BNA5* and *BNA6* were significantly under-expressed even in the absence of nicotinamide treatment. In contrast, *BNA7* was not induced by nicotinamide and was unaffected in *spp1Δ* cells (Figure 3 – figure supplement 1C). These results matched the RNAseq data for log and 48 hour-aged cells remarkably well (Figure 3C,D and figure supplement 1A,B,C compare left and right panels), showing that the impact of H3K4me3 is not restricted to ageing cells and therefore that H3K4me3 is required for normal expression and induction of the *BNA* genes.

The *BNA* genes are also induced to a lesser extent in *swd1Δ* mutants but the effect is less clear-cut as basal expression is more variable, ranging from much higher than wild type in *BNA4* to lower than wild type in *BNA6* (Figure 3C,D and Figure 3 – figure supplement 1A,B). The basal expression level of *BNA7* is also reduced in *swd1Δ* mutants and is not induced back to wild-type levels by nicotinamide (Figure 3 – figure supplement 1C). Again, these observations closely paralleled the ageing RNAseq data at 48 hours, and show that while loss of H3K4me3 reduces the mRNA level of multiple NAD+ biosynthesis genes, H3K4me1/2 has more complex gene specific effects.

COMPASS has at least one non-histone substrate (Zhang et al., 2005), so to ensure that the effects on NAD+ biosynthesis genes result from H3K4 methylation we performed further experiments in histone mutants H3K4A and H3K4R. As before we used nicotinamide to induce expression of the *BNA* genes in haploid cells growing at log phase in YPD, and observed that the H3K4 mutants largely phenocopied the COMPASS mutants; both mutants acted like *spp1Δ* for *BNA1, BNA5* and *BNA6* but like *swd1Δ* for *BNA4* and *BNA7* (Figure 3E,F and Figure 3 – figure supplement 1D-F). This is consistent with COMPASS acting through H3K4 methylation to alter the expression of the *BNA* genes, albeit through more than one mechanism (see discussion).

Together, these data show that COMPASS mutation affects the controlled expression of NAD+ biosynthesis genes directly via H3K4 methylation, rather than through indirect age-induced changes in metabolism. They further provide strong evidence for a critical role of H3K4me3 in the expression of yeast genes. Although the 2-fold under-expression observed here in *spp1Δ* mutants is not a huge effect, *BNA* gene mRNA levels are tightly auto-regulated in response to NAD+ levels so this is likely to limit the production of NAD+ for the lifetime of the organism.

### Age-linked changes in H3K4me3 distribution

Ageing in yeast is accompanied by widespread gene expression changes and a gradual reduction in histone density (Feser et al., 2010); western blot analysis of ageing MEP cells confirmed the progressive loss of histone H3 and revealed a similar decrease in H3K4me3 with age (Figure 4A). To understand how these differences affect individual promoters requires the genome-wide distribution of H3K4me3 to be determined, however mapping histone modifications in ageing yeast cells has proved extremely challenging as standard yeast chromatin immunoprecipitation (ChIP) protocols require 100-1000x more cells than are available from routine ageing cell purifications (explaining the very limited genome-wide ageing ChIP data available for yeast (Hu et al., 2014; Sen et al., 2015)). To facilitate the analysis of ageing chromatin, we optimised a low-cell number yeast ChIP protocol that yields ChIPseq libraries from the 10^6^ cells available through our standard ageing cell preparation methods. We used this protocol to obtain ChIPseq datasets for H3 and H3K4me3 in log phase cells and cells aged for 7.5, 24 and 48 hours (triplicate for log and 24 hours, duplicate for 7.5 and 48 hours, out of which only one 48 hour H3 sample failed quality control), along with matched input DNA. Input samples confirmed known age-linked genetic changes including amplification of rDNA and telomeres (Sinclair and Guarente, 1997), amplification of Chr. XII distal to the rDNA, and amplification of Ty elements (predicted given increase in Ty element RNAs)(Hu et al., 2014) (Figure 4 – figure supplement 1A), and all of these regions were therefore excluded from the chromatin analysis (Table S1). However, detection of these differences confirmed that ChIP samples derived from highly aged cells rather than contaminating daughters.

**Figure 4:**
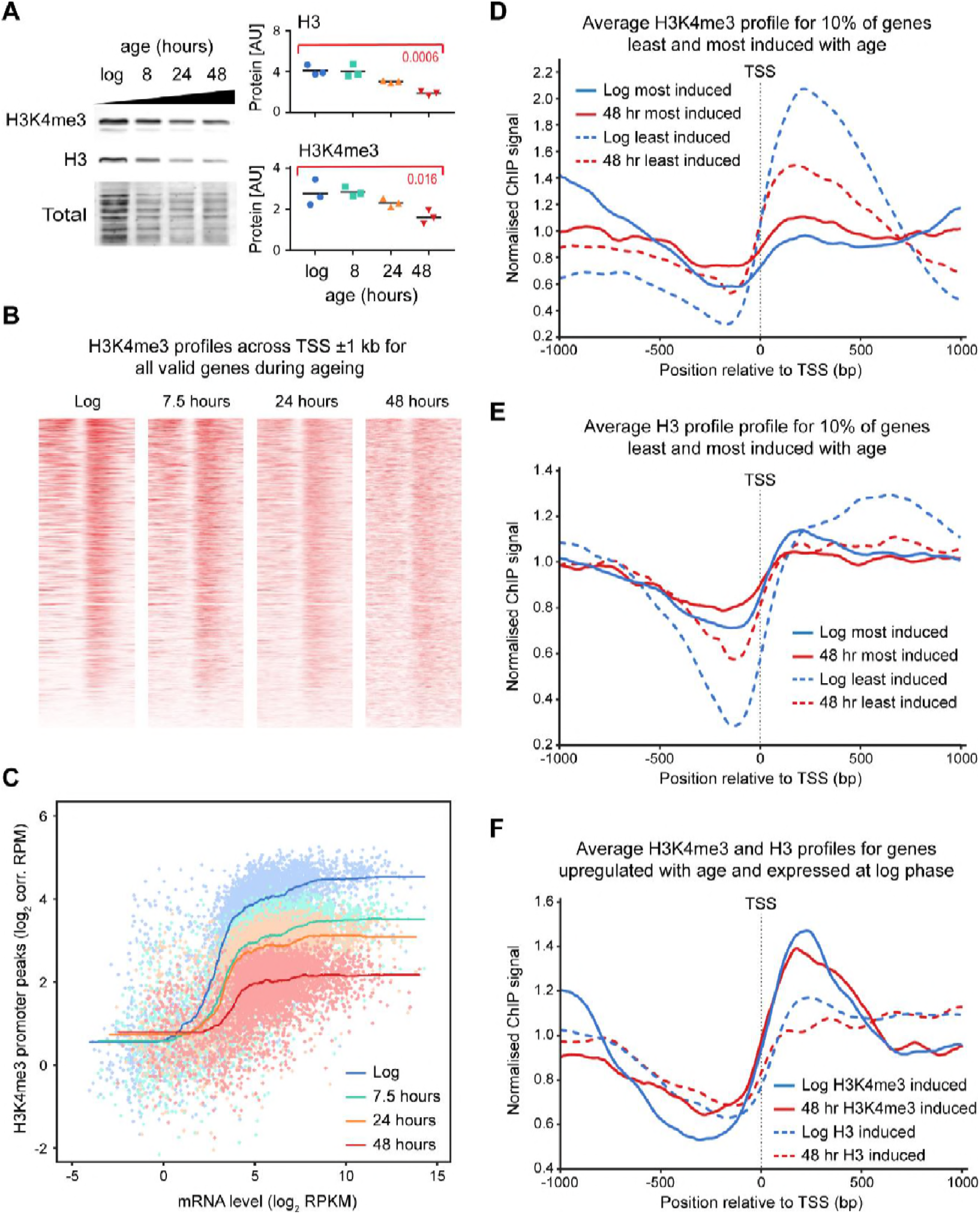
H3K4me3 declines with age despite genome-wide gene induction. **A**: Western blot analysis of histone H3 and H3K4me3 in cells aged for 0, 7.5, 24 or 48 hours. Signals were normalised to total protein and compared by one-way ANOVA, n=3, specific comparisons and p-values are shown in red. **B**: H3K4me3 profile at all transcriptional start sites (TSS) ±1 kb in wild-type cells across ageing. Genes that undergo age-linked copy number variation have been excluded. Intensity of colour corresponds to signal normalised to total valid reads per sample. **C**: Plot of H3K4me3 promoter peak versus gene expression at different ages. ChIP read counts were determined for 0-500 bp downstream of the TSS and normalised to background (see Materials and Methods). Each dot represents one gene. Lines represent smoothed data. **D**: Average profile of H3K4me3 ChIP signal for 1 kb either side of the transcriptional start site, average of all samples per time point (n=3 for log and 24 hours, n=2 for 7.5 and 48 hours), showing the 10% of genes most (481) and least induced (482) with age. **E**: As D, showing H3 ChIP signal. **F**: Average profile of H3 and H3K4me3 ChIP signals for 1 kb either side of the transcriptional start site as in D, showing genes (151) with normalised RPKM>3 in log phase and >2 fold induction from log to 48 hours based on RNAseq data.

We first examined the H3K4me3 peak associated with all annotated promoters, and as expected based on the western blot analysis, this peak progressively decreased in prominence with age while noise noticeably increased (Figure 4B, Figure 4 – figure supplement 1B). ChIPseq data does not maintain information about absolute quantities of histone bound DNA, only relative concentrations at different sites, so we suspected that the decline in the observed H3K4me3 peak stemmed from a decrease in signal relative to noise. However, ChIP is prone to technical variation in enrichment (signal to noise ratio). To gauge whether the observed decrease in signal was technical or biological, we calculated ratios of the 10^th^ percentile value (noise) to the 90^th^ percentile value (signal) (Figure 4 – figure supplement 1C). Despite some variation in enrichment, these ratios decreased across time points in the H3K4me3 samples confirming that the H3K4me3 peak is progressively lost with age. In contrast, the H3 data remained largely stable with age, as H3 is not specifically enriched in particular regions of the genome. The measured nucleosome free region (NFR) does become slightly reduced with age, a result of the histone density across the genome being lower and therefore the difference in signal between the NFR and surrounding chromatin being smaller (Figure 4 – figure supplement 1D).

While these results are in accord with existing literature and our western blotting data, they lead to a surprising conclusion: since gene expression generally increases with age while the H3K4me3 peak is lost, the normal positive correlation between H3K4me3 and gene expression must decrease substantially with age. Comparing mRNAseq to H3K4me3 signals at the different ages confirms that this is the case (Figure 4C), with the difference in average H3K4me3 peak signal between low and highly expressed genes being ~16-fold in log phase but ~2-fold in 48 hour-aged cells.

We then examined the chromatin states of promoters in sets of genes that show characteristic age-linked behaviours. Firstly, we compared the 10% of genes least and most induced with age: the least induced genes show a clear loss of H3K4me3 with time as expected, but more surprisingly the deposition of H3K4me3 at the most induced genes is minimal and no defined promoter associated H3K4me3 peak forms for genes in this category (Figure 4D). Furthermore, the already minimal NFR of the most induced genes becomes even less defined with age (Figure 4E). As previously noted, the genes most induced with age are very poorly expressed at log phase, so key promoter-associated factors may not be present (Hu et al., 2014). We therefore also examined genes induced 2-fold or more with age and detectably expressed (normalised RPKM>3) at log phase; in this set, both the H3K4me3 peak declines with age and the NFR becomes less distinct (Figure 4F).

Overall, our data show that promoter H3K4me3 declines with age such that in very old cells this mark becomes a poor predictor of gene expression, even on genes that are robustly induced with increasing age.

## Discussion

The predicted role of H3K4me3 in facilitating transcription has proved challenging to validate. Here we have shown that a large subset of yeast genes depend on H3K4me3 for full induction. This effect becomes prevalent during the ageing process in which huge numbers of genes are induced coincident with a decline in the repressive capacity of H3K4 methylation. Our results provide a clear demonstration that H3K4me3 is indeed an epigenetic mark that facilitates gene expression, and one that plays a critical role in maintaining cell viability across the life-course.

### Gene expression outcomes of H3K4 methylation

H3K4 methylation has a remarkably complex effect on gene expression. Transcriptomic approaches have detected repressive functions (Guillemette et al., 2011; Jaiswal et al., 2017; Lenstra et al., 2011; Margaritis et al., 2012; Venkatasubrahmanyam et al., 2007), although in most cases this can probably be attributed to H3K4me2, with the possible exception of ribosomal protein genes (Weiner et al., 2012). Evidence that H3K4me3 can promote gene expression in yeast is rarer; this was reported in early studies but has not been effectively reproduced (Miller et al., 2001; Santos-Rosa et al., 2002), see (Margaritis et al., 2012) for a discussion of technical reasons for this. Setting aside technical issues, if H3K4 methylation has both repressive and inducing activities, the outcome of COMPASS mutations on individual genes would be expected to vary by gene, by genetic background and by experimental system, making broad conclusions rather treacherous.

Antagonism between activating and repressive activities would help explain the conflicting data in the literature, and close examination of the data in Figure 3 provides support for this: loss of Spp1 reduces expression for most of the *BNA* genes, whereas loss of Swd1 has a milder and more variable effect that likely results from the summed outcomes of repressive H3K4me2 and activating H3K4me3 (see summary of H3K4me3 and H3K4me2 effects on the *BNA* genes in Figure 5). However for other genes, the balance and direction of these effects is clearly different, for example at *BNA7* H3K4me2 appears to be activating while H3K4me3 has no effect. Therefore, the impact of H3K4 methylation cannot be generalised but must be considered on a gene-by-gene basis taking into account both repressive and activating effects.

**Figure 5:**
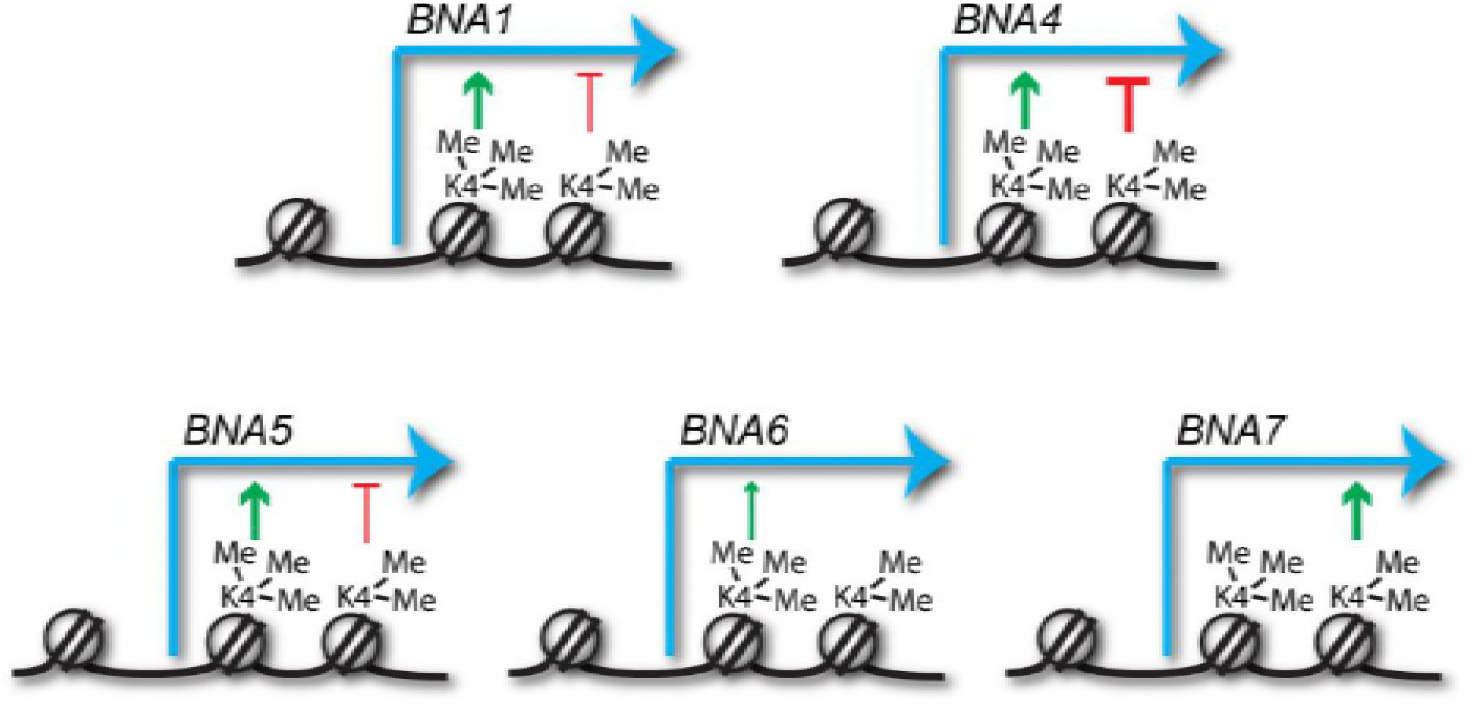
Predicted impacts of H3K4me2 and H3K4me3 on *BNA* gene expression. Schematic of the different impacts of H3K4me2 and H3K4me3 on expression of the *BNA* genes analysed in Figure 3 and Figure 3 – figure supplement 1. Depiction of these modifications on the +2 and +1 nucleosomes respectively is for illustrative purposes only and simply indicates that H3K4me2 normally spreads further into the gene body than H3K4me3.

How might H3K4me3 promote gene expression? Multiple H3K4me3-binding proteins are present in yeast, which in turn are members of histone modifying and repositioning complexes. Binding of Yng1, Yng2 or Sgf29 would recruit NuA3, NuA4 or SAGA histone acetyl transferase complexes respectively, promoting local histone acetylation that should enhance transcriptional activation (Agalioti et al., 2002; Bian et al., 2011; Pokholok et al., 2005; Steunou et al., 2016; Taverna et al., 2006). Loss of recruitment of NuA3 and NuA4 to H3K4me3 has remarkably little effect on gene expression under normal conditions (Choy et al., 2001; Taverna et al., 2006), but may well impact a specific subset of genes when induced or during dramatic chromatin remodelling of the sort that occurs during ageing; indeed delays in induction of individual genes have been reported in mutants lacking H3K4me3-linked acetyltransferase recruitment (Bian et al., 2011; Morillon et al., 2005).

### The impact of H3K4 methylation on ageing cells

Our experiments very clearly show that H3K4 methylation and in particular H3K4me3 is critical for the maintenance of a normal lifespan. We observed 43% of yeast genes to be differentially expressed during ageing, of which 17% show significantly different expression levels in COMPASS mutants after 48 hours. This means that as cells age, loss of H3K4 methylation has a dramatic effect on maintenance of normal gene expression patterns. We observed a progressive loss of the repressive capacity of H3K4 methylation as cells age, which likely relates to the age-linked reduction in histone levels; Margaritis *et al.* observed that *hht1Δ hhf1Δ* mutants show a similar pattern of gene upregulation to *set1Δ* cells (Margaritis et al., 2012). However, this is probably not the cause of the lifespan defect, as this repressive activity is not impacted by *spp1Δ* whereas lifespan is similar in *swd1Δ* and *spp1Δ* cells. Rather, the most prominent gene expression effect was a failure to fully induce hundreds of genes in COMPASS mutants including *spp1* Δ. We validated this role of COMPASS for expression of NAD+ biosynthesis genes, but these constitute only a handful of the significantly mis-expressed genes and are unlikely to explain the shortened lifespan of COMPASS mutants alone. In reality, given the widespread age-linked gene mis-regulation in COMPASS mutants, it is not surprising that the lifespan of these cells is profoundly reduced.

Unexpectedly, we observed that the widespread transcriptional induction which accompanies ageing does not entail an increase in H3K4me3, such that the correlation between gene expression and H3K4me3 progressively declines. This was surprising for two reasons: firstly, the effect of COMPASS mutation is much stronger in highly aged cells despite the H3K4me3 mark being at its weakest. Our analysis shows that gene expression defects in COMPASS mutants are progressive starting from log phase, and we interpret the underexpression at 48 hours as the cumulative outcome of defective gene expression throughout life, which would be little affected by the low H3K4me3 in aged cells. Secondly, as described above recruitment of COMPASS involves the Paf complex, Rad6/Bre1, H2B, Ser-5 phosphorylated RNA pol II CTD and nascent mRNA. As gene expression is induced with age, we would expect the latter two elements to rise and increase COMPASS recruitment, raising local H3K4me3 irrespective of declining histone occupancy. Hu *et al* noted that many of the genes induced during ageing lack an NFR (Hu et al., 2014); in other words, these may not act as classical promoters and therefore may not have promoter-associated Paf and/or Rad6/Bre1 to recruit COMPASS. However, many genes induced with age are already expressed in young cells and therefore possess an active promoter and defined NFR (see Figure 4F). Although the age-linked increase in mRNA levels could also stem from post-transcriptional regulation or passive mRNA accumulation, this is unlikely given that many of the changes are recapitulated by H3 depletion and impaired in COMPASS mutants, both of which would be most likely to affect transcription. It seems therefore, that parts of the transcriptional machinery which normally recruit COMPASS, and others that normally respond to H3K4me3, become less active with age and discovering the nature of these defects will provide an important focus for future studies into age-linked gene expression change.

## Materials and Methods

### Strains and media

Yeast strains were constructed by standard methods and are listed in Table S2, oligonucleotide sequences are given in Table S3 and plasmids given in Table S4. To construct *set1^H1017L^* we used the break-mediated *Delitto Perfetto* system (Storici and Resnick, 2006): after integration of the CORE cassette, *Sce*I expression was induced using 2% galactose and oligonucleotides OCC339/40 transformed in followed by selection for fluoroorotic acid and against G418. All cells were grown in YPD media (2% peptone, 1% yeast extract, 2% glucose) at 30°C with shaking at 200 rpm. Media components were purchased from Formedium and media was sterilised by filtration. For MEP experiments, cells were inoculated in 4ml YPD and grown for 6-8 hours then diluted in 25ml YPD and grown for 16-18 hours to 0.2-0.6x10^7^ cells/ml. If required, cells were labelled (see below), then cells were inoculated in YPD at 2x10^4^ cells/ml containing 5μg/ml ampicillin (Melford A0104) and 1μM β-estradiol (Sigma E2758). For viability assay time points, 10μl of culture was diluted with 40μl water and spread on a YPD plate.

### MEP cell labelling and purification

For biotin labelling, 0.25x10^7^ cells per sample were harvested by centrifugation (15 s at 13,000 g), washed twice with 125μl PBS and re-suspended in 125μl of PBS containing ~3mg/ml Biotin-NHS (Sigma B1022 or Pierce 10538723). Cells were incubated for 30 min on a wheel at room temperature, washed once with 125μl PBS and re-suspended in YPD. For 7.5 hour time points, cells were inoculated in 15ml YPD + 5μg/ml ampicillin and grown for 2 hours before addition of 1μM β-estradiol (Sigma E2758). For 24 and 48 hr time points, cells were directly inoculated in 125ml YPD containing 5μg/ml ampicillin and 1mM β-estradiol. For 24/48 hour time points, no contamination of labelled non-aged cells was observed by bud scar staining as these cells presumably lyse, and this strategy helped keep the culture density low (cultures did not exceed 0.75x10^7^ cells/ml in 48 hours). Cells were harvested by centrifugation for 1 min, 4600 rpm in 15/50 ml tubes and immediately fixed by resuspension in 70% ethanol and storage at −80°C. Alternatively, formaldehyde (Thermo 11586711) was added to 1% and cells incubated at room temperature for 15 min before quenching with 125 mM glycine, harvested by centrifugation as above and washed three times with PBS before freezing on N2 and storage at −80°.

Rapid purification for RNA or protein: Percoll gradients (1 for 7.5 hour or 2 for 24/48 hour time points) were formed by vortexing 1ml Percoll (Sigma P1644) with 110μl 10x PBS in 2ml tubes and centrifuging 15 min at 15,000 g, 4 °C. Ethanol fixed cells were defrosted and washed 1x with 1 volume of cold PBSE (PBS + 2 mM EDTA) before resuspension in ~250 μl cold PBSE per gradient and layering on top of the pre-formed gradients. Gradients were centrifuged for 4 min at 2,000 g, then the upper phase and brown layer of cell debris removed and discarded. 1 ml PBSE was added, mixed by inversion and centrifuged 1 min at 2,000 g to pellet the cells, which were then re-suspended in 1 ml PBSE per time point (re-uniting samples split across two gradients). Cells were briefly sonicated in a chilled Bioruptor 30 s on low power, then 25 μl Streptavidin microbeads added (Miltenyi Biotech 1010007) and cells incubated for 5 min on a wheel at room temperature. Meanwhile, 1 LS column per sample (Miltenyi Biotech 1050236) was equilibrated with cold PBSE in 4 °C room. Cells were loaded on columns and allowed to flow through under gravity, washed with 2 ml cold PBSE and eluted with 1 ml PBSE using plunger. Cells were re-loaded on the same columns after re-equilibration with ~500 μl PBSE, washed and re-eluted, and this process repeated for a total of three successive purifications. 50 μl cells were set aside for quality control, while the remainder were pelleted by centrifugation and processed directly for RNA or protein.

For quality control, the 50 μl cells were diluted to 300 μl final volume containing 0.3% triton X-100, 0.3 μl 1mg/ml streptavidin 594 (Life Technologies S11227), 0.6 μl 1 mg/ml WGA-488 (Life Technologies W11261) and DAPI. Cells were stained for 15 min at room temperature in the dark, washed once with PBS then re-suspended in 5 μl VectaShield (Vectorlabs H-1000). Purifications routinely yielded 80-90% streptavidin positive cells with appropriate bud-scar numbers.

Purification of formaldehyde-fixed cells for ChIP: Percoll gradients (1 for 7.5 hour or 2 for 24/48 hour time points) were formed by vortexing 1.16ml Percoll (Sigma P1644) with 42 μl 5 M NaCl, 98 μl water in 2 ml tubes and centrifuging 15 min at 15,000 g, 4 °C. Cells were defrosted on ice, re-suspended in ~250 μl cold PBSE per gradient and layered on the preformed gradients. Gradients were centrifuged for 20 min at 1,000 g, then the upper phase and brown layer of cell debris removed and discarded. 1 ml PBSE was added, mixed by inversion and centrifuged 1 min at 2,000 g to pellet the cells, which were then re-suspended in 1ml PBSE per time point (re-uniting samples split across two gradients). 25 μl Streptavidin magnetic beads were added (Miltenyi Biotech 1010007) and cells incubated for 30 min on a wheel at room temperature. Meanwhile, 1 LS column per sample (Miltenyi Biotech 1050236) was equilibrated with cold PBSE in 4 °C room. Cells were loaded on columns and allowed to flow through under gravity, washed with 8 ml cold PBSE and eluted with 1ml PBSE using plunger. Cells were re-loaded on the same columns after re-equilibration with ~500 μl PBSE, washed and re-eluted, and this process repeated for a total of three successive purifications. Final elution was to 1.5 ml tubes (Treff 1130153). 50 μl cells were set aside for quality control, while the remainder were pelleted by centrifugation and processed directly for ChIP.

### Low cell number RNA purification and library preparation

Cells were re-suspended in 50 μl Lysis/Binding Buffer (from mirVANA kit, Life Technologies AM1560), and 50 μl 0.5 μm zirconium beads (Thistle Scientific 11079105Z) added. Cells were lysed with 5 cycles of 30 s 6500 ms^−1^ / 30 s ice in an MP Fastprep bead beater, then 250 μl Lysis buffer added followed by 15 μl miRNA Homogenate Additive and cells were briefly vortexed before incubating for 10 minutes on ice. 300 μl acid phenol: chloroform was added, vortexed and centrifuged 5 min at 13,000 g, room temperature before extraction of the upper phase. 400 μl room temperature ethanol and 2 μl glycogen (Sigma G1767) were added and mixture incubated for 1 hour at −30 °C before centrifugation for 15 minutes at 13,000 g, 4 °C. Pellet was washed with cold 70% ethanol and re-suspended in 10 μl water. 1 μl RNA was glyoxylated and analysed on a BPTE mini-gel as described (Sambrook and Russell, 2001), and 0.2 μl quantified using a PicoGreen RNA kit (Life Technologies R11490).

500 ng RNA was used to prepare libraries using the NEBNext Ultra Directional mRNAseq kit (NEB E7420, E7490, E7335, E7500, E7710) as described with modifications: Reagent volumes for RT, second strand synthesis, tailing and ligation were reduced by 50%, while libraries were amplified for 12 cycles using 2 μl each primer per reaction before two rounds of AMPure bead purification as described in the kit at 0.9x ratio prior to analysis by quality control using a Bioanalyzer.

### Low cell number ChIP and library preparation

Cells were re-suspended in 50 μl cold CLB (50mM HEPES pH 7.0, 14mM NaCl, 1mM EDTA, 1% Triton X100, 0.1% sodium deoxycholate, 0.1% SDS, cOmplete Protease Inhibitors (Roche 1836170)) with 50 μl zirconium beads and lysed with 5 cycles of 30 s 6500 ms^−1^ / 30 s ice in an MP Fastprep bead beater then decanted to a Covaris 520045 microTUBE and topped up to 130 μl with CLB. Sonication was performed on a Covaris 220: Duty Factor 5 %, PIP 130 W, 900s, 200 cycles per burst, 8.9 °C. Samples were decanted to a 1.5 ml tube and Covaris tube washed with CLB and combined to give 180 μl total volume, which was centrifuged 5 min top speed at 4 °C and the pellet discarded. Lysate was pre-cleared by incubation with 10 μl of preequilibrated Protein-G magnetic beads (Life Technologies 10004D), then 30 μl set aside for total DNA extraction and 50 μl used per immunoprecipitation (IP). Antibodies were added at 1:50 dilution (rabbit anti-H3, CST 2650 or rabbit anti-H3K4me3, CST 9751), one IP was processed without antibody, and IPs were incubated over night at 4 °C on a wheel.

10 μl pre-equilibrated protein-G beads in 25 μl CLB were added to each IP and incubated 2-3 hrs at 4 °C on a wheel. Beads were then washed once each at room temperature for 5 minutes on a wheel with: CLB, CLB with 0.5 M NaCl, wash buffer (10 mM Tris pH 8.0, 0.25 M LiCl, 0.5% NP-40, 0.5% sodium deoxycholate, 1 mM EDTA), TE, then eluted overnight at 65 °C with 200 μl elution buffer (50 mM Tris/HCl pH 8.0, 10 mM EDTA, 1% SDS). Input samples were diluted to 200 μl with TE, treated with 1 μl 1 mg/ml RNase A at 37 °C for 1 hour then incubated overnight at 65 °C. Input and IP samples were treated with 1 μl proteinase K for 2 hours at 55 °C, then purified by phenol:chloroform extraction, precipitated and re-suspended in 5 μl TE.

After quality control of 10% by qPCR, libraries were synthesised using a NEBNext DNA Ultra II kit (NEB E7645) with modifications: Concentration of DNA was increased by reducing the volumes of the end repair and ligation steps 5-fold, with reagent quantities reduced in kind. After clean-up with 0.9x AMPure beads, PCR was performed in 50 μl volumes according to manufacturer’s instructions. Libraries were purified with 0.9x AMPure beads and eluted in 30μl, then a size selection was performed by incubating first with 18μl AMPure beads, discarding the beads then purifying the remaining DNA from the supernatant with an additional 9 μl AMPure beads.

### Nicotinamide treatment, RNA purification and northern analysis

Cells were grown in 4 ml YPD for 6 hours, diluted and grown overnight in 25 ml YPD to 0.2-0.5x10^7^ cells/ml. Cells were diluted to 0.06x10^7^ cells/ml in 12.5 ml YPD and grown for 1.5 hours before addition of 12.5 ml YPD with or without 2 mM nicotinamide (Sigma N3376) and grown for 6 hours before harvesting 2x10^7^ cells by centrifugation and freezing on N2. RNA was extracted by the GTC phenol miniprep method, and 1.5 μl of 6 μl eluate run on 1% BPTE gels, imaged and blotted as described (Cruz and Houseley, 2014). Gels were probed for *BNA4* and *BNA6* then re-probed for *BNA1*, or for *BNA5* and *BNA7*, in UltraHyb (Life Technologies AM8670) using RNA probes listed in Table S5 as described (Cruz and Houseley, 2014).

### Protein purification and western blotting

Proteins were purified, separated and blotted as described with modifications (Frenk et al., 2014): aged MEP cells were re-suspended in Laemmli buffer containing 100mM DTT and separated on 15% acrylamide gels, and Odyssey blocking buffer PBS (LI-COR 927-40000) was used for antibody hybridisations. Membranes were probed with antibodies listed in Table S6 at given dilutions before imaging on a LI-COR Odyssey 9120 imager. Total protein detection was performed by staining membranes with REVERT Total Protein Stain solution (LiCor 926-11010) after antibody detection.

### Image processing and data analysis

Northern blots images were quantified using ImageQuant v7.0 (GE), western blots using Aida v3.27.001 (Fuji). Images for publication were processed using ImageJ v1.50i, by cropping and applying minimal contrast enhancement. Statistical analysis of viability scores and northern/western data was performed using GraphPad Prism v7.03.

### Sequencing and bioinformatics

After adapter and quality trimming using Trim Galore (v0.4.2), RNAseq data was mapped to yeast genome R64-2-1 using HISAT v2.0.5 (Kim et al., 2015). Mapped data was imported into SeqMonk v1.39.0 (https://www.bioinformatics.babraham.ac.uk/projects/seqmonk/) and normalised based on the total number of reads mapping to the antisense strand of annotated open reading frames (opposite strand specific libraries), excluding the rDNA locus and the mtDNA. Exclusion of rDNA mapping reads from the normalisation is particularly important as in ageing cells a substantial fraction of reads derive from polyadenylated rDNA ncRNAs and can skew the distribution. DESeq2 analyses (Love et al., 2014) was performed within SeqMonk, comparing either five replicates of different time points in the wild-type dataset or comparing the five replicates of wild-type to pooled data from *swd1* Δ, *set1* and *spp1Δ* mutants at a given time point, using a p-value cut-off of 0.01 for analyses except comparisons of wild-type versus *spp1Δ* where a cut-off of 0.05 was applied. Hierarchical clustering analysis was performed using SeqMonk, and GO analysis of individual clusters performed using GOrilla (http://cbl-gorilla.cs.technion.ac.il/) (Eden et al., 2007; Eden et al., 2009). TSS data was obtained from (Pelechano et al., 2013) – the strongest TSS in GLU or GAL was used where TSS data was available, otherwise the start of the CDS was substituted.

After adapter and quality trimming using Trim Galore (v0.4.2), ChIPseq data was mapped to yeast genome R64-2-1 using Bowtie 2 (v2.3.2). Mapped data was analysed using SeqMonk (v1.39.0, https://www.bioinformatics.babraham.ac.uk/projects/seqmonk/). Reads mapping to Excluded Regions (genomically amplified with age, see Table S1, Figure 4 – figure supplement 1A) were excluded from the analysis. Because of different global enrichment during aging, samples were normalised to background signal (Fig. 4C). Read counts of 500 bp tiles across the genome were determined, and the value at the 10^th^ percentile used to define background signal. ChIP enrichment was calculated over the background of the sample. For profile plots (Fig. 4D-F) the enrichment/depletion normalised to total count of valid reads is shown.

All raw mRNAseq and ChIPseq data has been deposited at GEO under accession number GSE107744.

## Acknowledgements

We thank Kristina Tabbada and Clare Murnane in the BI Next Generation Sequencing Facility for data generation, Felix Krueger, Anne Segonds Pichon and Simon Andrews of the BI Bioinformatics facility along with Mikhail Spivakov for statistical and bioinformatics advice, Gavin Kelsey for critical reading, and Dan Gottschling for the MEP strains. Funding was from the Wellcome Trust [088335,110216], the BBSRC [BI Epigenetics ISP: BBS/E/B/000C0423], and the ERC [EpiGeneSys Network].

---

**Figure 1 – figure supplement 1:**
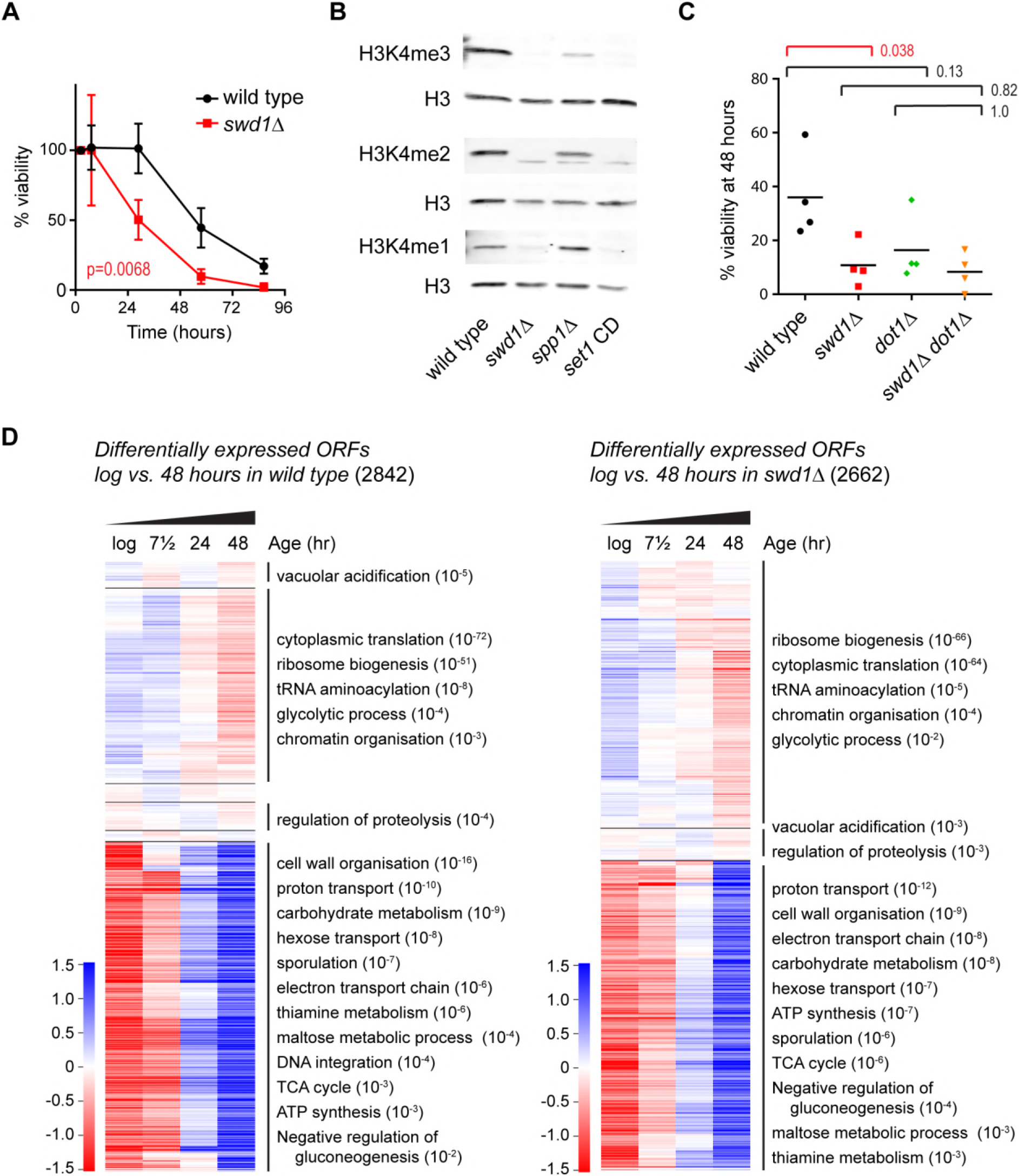
Supplement to the ageing transcriptome in wild type and COMPASS mutants. **A**: Viability of mother cells across 72 hours in liquid YPD culture using the MEP system for wild type and *swd1Δ* mutants. p-value calculated based on area under curve n=5 for wild type, n=3 for *swd1* Δ. **B**: Western blot analysis of H3K4me3, H3K4me2 and H3K4me1 in log phase MEP cells. Three replicate membranes were probed with mouse anti-H3 and rabbit anti-H3K4me1/2/3 using a 2-colour detection system, such that each methyl-H3K4 antibody has a matched H3 control. **C**: Viability of mother cells after 48 hours in liquid YPD culture using the MEP system. p-values calculated by one-way ANOVA, n=4. **D**: Hierarchical clustering analysis of log2-transformed protein-coding mRNA change for genes differentially expressed between log phase and 48 hours ageing in wild-type (left) and in *swd1*Δ (right), assessed using DESeq2 p<0.01 n=5 per time point (wild type) or n=4 per time point (swd1Δ). Each annotated cluster shows significant GO enrichments assigned by GOrilla, FDR-corrected p-value<0.05. GO terms representing similar functions have been collapsed; full GO analyses are presented in Supplementary Material.

**Figure 1 – figure supplement 2:**
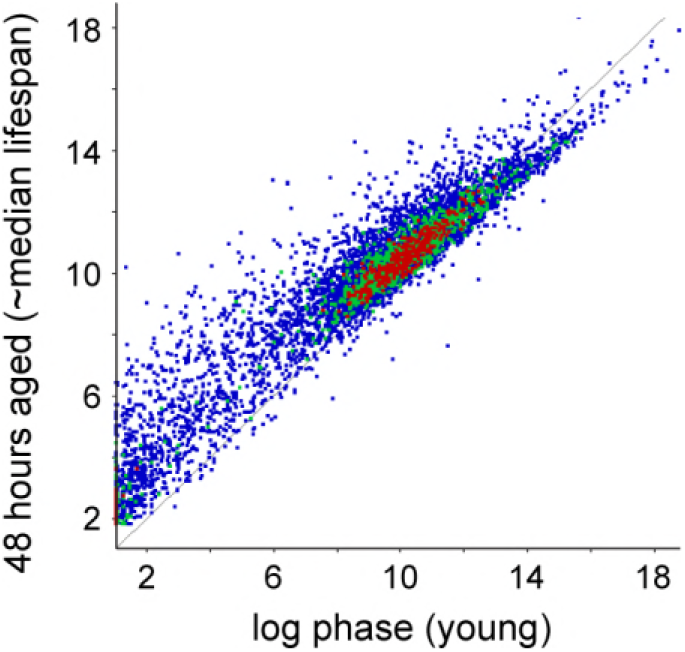
Age-linked change in gene expression profile. Scatter plot of log2-transformed mRNAseq read counts from protein coding genes normalised for ORF length comparing log phase cells to 48 hour-aged cells. Data averaged across 5 replicates per condition.

**Figure 2 – figure supplement 1:**
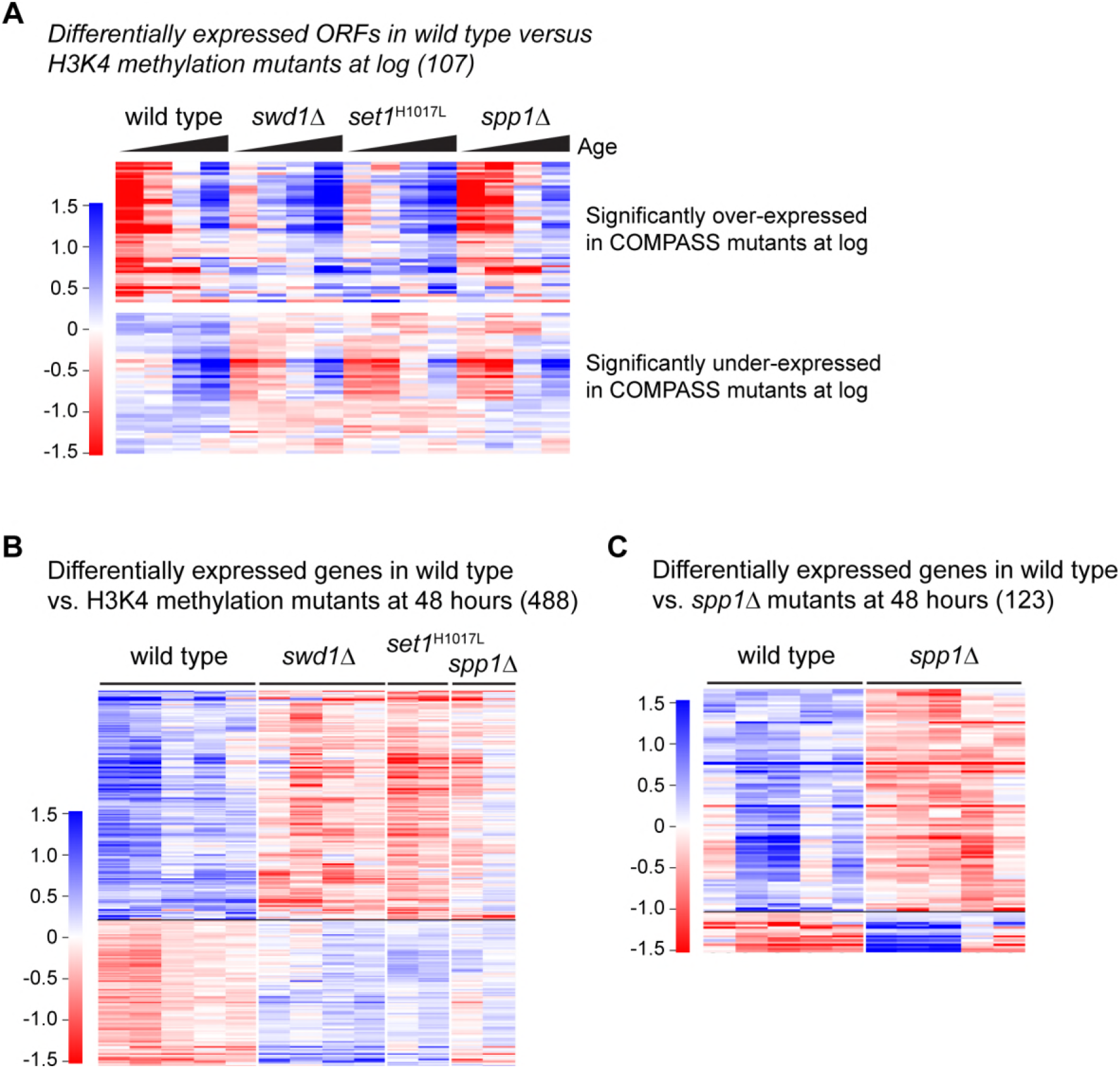
Supplement to COMPASS is required for gene expression in ageing cells. **A**: Hierarchical clustering analysis of log2-transformed protein-coding mRNA levels for genes significantly differentially expressed between wild type and COMPASS mutants at 48 hours ageing (same set of genes as Figure 2B). All 48 hour replicates are shown to demonstrate the reproducibility of gene expression differences. **B**: Analysis as in A showing reproducibility across 5 replicates of gene expression differences between aged wild-type and *spp1Δ* cells at 48 hours, gene set as in Figure 2C.

**Figure 3 – figure supplement 1:**
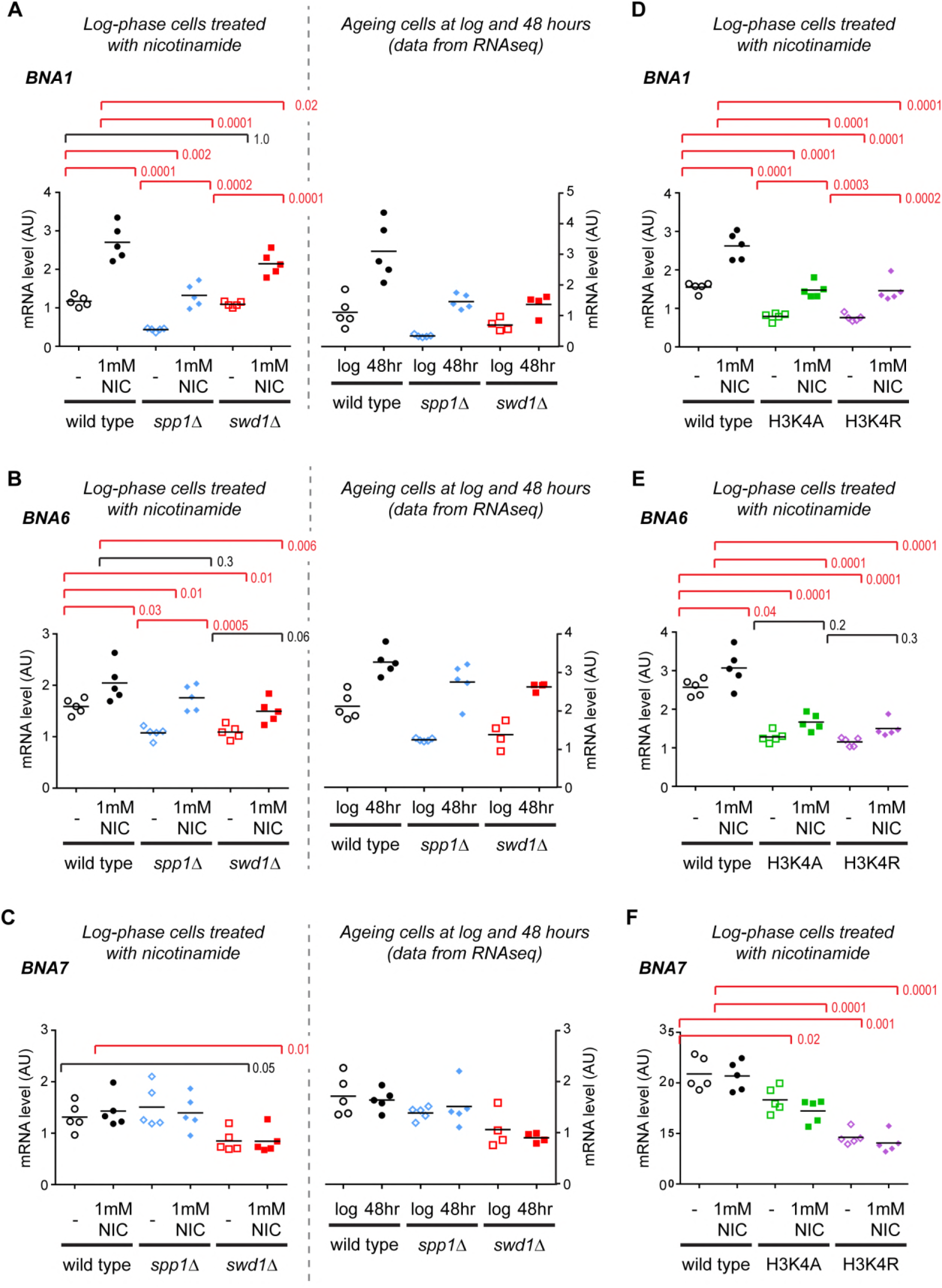
Supplement to validation of gene expression differences in COMPASS mutants. **A-F**: Quantification of *BNA* gene induction by northern blot for *BNA1, BNA6* and *BNA7*, performed as in Figure 3C.

**Figure 4 – figure supplement 1:**
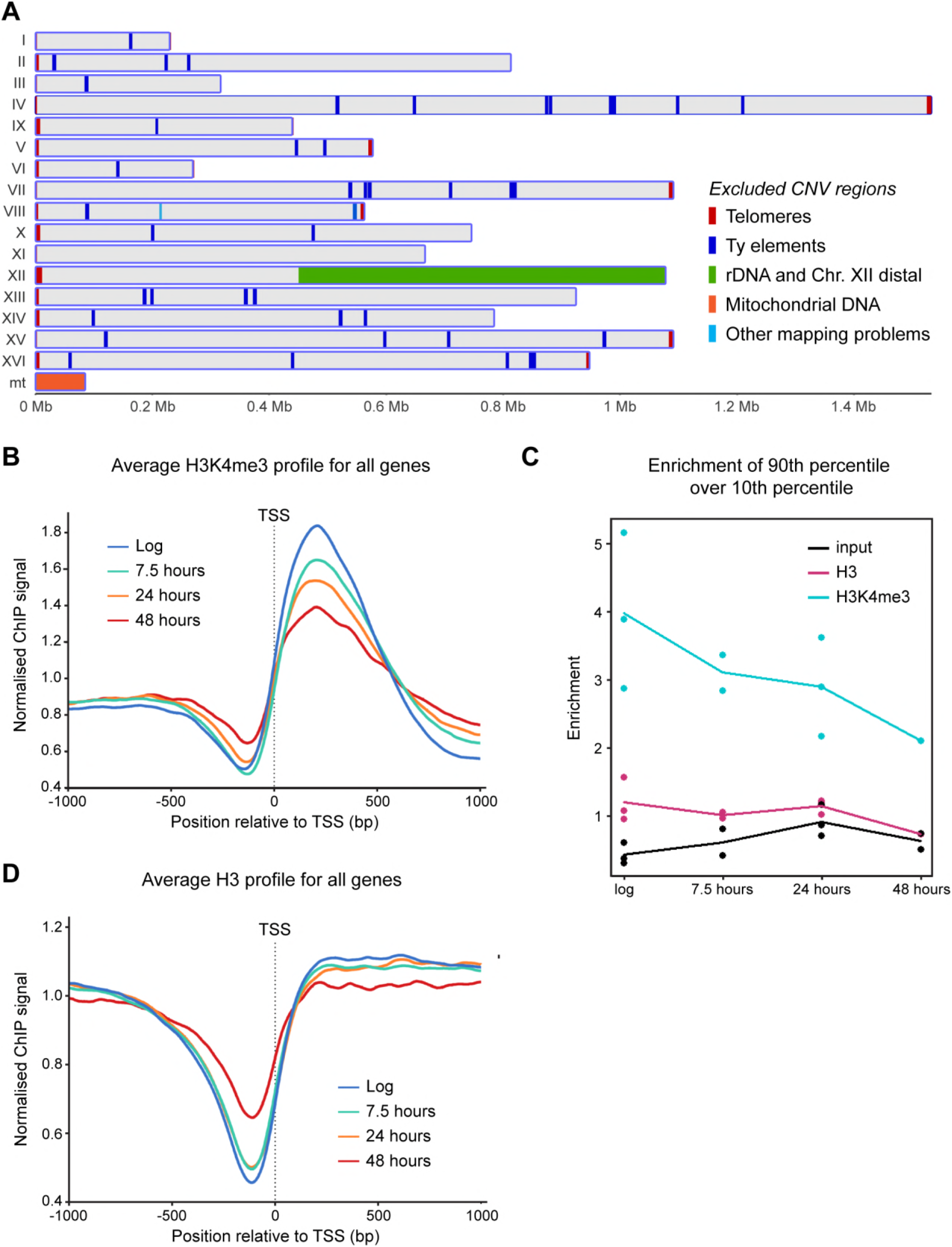
Supplement to H3K4me3 declines with age despite genome-wide gene induction. **A**: Genomic regions subject to age-linked CNV detected by comparison between input samples; these were excluded from ChIP analysis. **B**: Average profile of H3K4me3 ChIP signal for 1 kb either side of the transcriptional start site, average of all samples per time point (n=3 for log and 24 hours, n=2 for 7.5 and 48 hours). Data is the same as in Figure 4B, this is an alternate representation. **C**: Log2 read count of the 90^th^ percentile over the 10^th^ percentile for each ChIP library, showing that signal to noise decreases with age for H3K4me3 but not for H3 or input. The decrease over time outweighs the technical variability. **D**: Average profile of H3 ChIP signal for 1 kb either side of the transcriptional start site, average of all samples per time point (n=3 for log and 24 hours, n=2 for 7.5 hours and n=1 for 48 hours).

